# Extracellular thiamine concentration influences thermogenic competency of differentiating neck area-derived human adipocytes

**DOI:** 10.1101/2023.04.13.536432

**Authors:** Boglárka Ágnes Vinnai, Rini Arianti, Ferenc Győry, László Fésüs, Endre Kristóf

## Abstract

Brown adipose tissue (BAT) dissipates energy in the form of heat majorly via the mitochondrial uncoupling protein 1 (UCP1). The activation of BAT, which is enriched in the neck area and contains brown and beige adipocytes in humans, was considered as a potential therapeutic target to treat obesity. Therefore, finding novel agents that can stimulate the differentiation and recruitment of brown or beige thermogenic adipocytes are important subjects for investigation. The current study investigated how the availability of extracellular thiamine (vitamin B1), an essential cofactor of mitochondrial enzyme complexes that catalyze key steps in the catabolism of nutrients, affects the expression of thermogenic marker genes and proteins and subsequent functional parameters during *ex vivo* adipocyte differentiation. Therefore, we differentiated primary human adipogenic progenitors that were cultivated from subcutaneous (SC) or deep neck (DN) adipose tissues in the presence of gradually increasing thiamine concentrations during their 14 day long differentiation program. Higher thiamine levels resulted in increased expression of thiamine transporter 1 and 2 both at mRNA and protein levels in human neck area-derived adipocytes. Gradually increasing concentrations of thiamine led to increased basal, cAMP-stimulated, and proton-leak respiration along with elevated mitochondrial biogenesis of the differentiated adipocytes. The extracellular thiamine availability during adipogenesis determined the expression levels of UCP1, PGC1a, CKMT2, and other browning-related genes and proteins in primary SC and DN-derived adipocytes in a concentration-dependent manner. Providing abundant amounts of thiamine further increased the thermogenic competency of the adipocytes. Our study raises the possibility of a novel strategy with long-term thiamine supplementation, which can enhance the thermogenic competency of differentiating neck area-derived adipocytes for preventing or combating obesity.

## 1 Introduction

Obesity is defined as the abnormal or excessive accumulation of fat, which has a long-term negative impact on health, as it contributes to the development of a number of chronic conditions and diseases [Panuganti *et al*., 2021], such as type 2 diabetes mellitus (DM), cardiovascular diseases, hypercholesterolemia, asthma, or certain type of cancers [Safei *et al*., 2021]. World Health Organization (WHO) reported that since 1975, the prevalence of obesity has almost tripled worldwide [WHO, 2021].

Two types of adipose tissue have been identified: white adipose tissue (WAT), which is majorly responsible for energy storage, and brown adipose tissue (BAT), which maintains core body temperature through non-shivering thermogenesis [Ikeda *et al*., 2018]. White adipocytes contain one large unilocular lipid droplet and few mitochondria, whereas brown adipocytes possess multilocular lipid droplets and large amounts of mitochondria that express uncoupling protein 1 (UCP1) which is referred as the major molecular marker of thermogenic fat cells [Dragoo *et al*., 2021; Cohen and Kajimura, 2021]. Another, inducible form of thermogenic adipocytes is called beige (also known as brite) adipocytes. Beige adipocytes possess a transitional phenotype between white and brown adipocytes. They are considered as the subset of WAT but have several properties of brown adipocytes, including increased UCP1 expression, multilocular triglyceride storage, mitochondria density as well as vascular supply, all of which promote the process of adaptive thermogenesis. The differentiation of beige adipocytes can be induced e.g. by cold, physical exercise, peroxisome proliferator-activated receptor (PPAR) γ, or ß3-receptor agonists which drive the process termed „browning” [Dragoo *et al*., 2021; Cohen and Kajimura, 2021].

Human BAT was mainly considered as a tissue type that can only be found in newborns in the axillary, cervical, perirenal, and periadrenal regions maintaining the proper core body temperature without shivering [Wu *et al*., 2013]. Because the amount of BAT decreases shortly after birth, it has been considered for a long period of time to be functionally insignificant in adults. However, several studies using positron emission tomography (PET) provided evidence that adults have significant amounts of BAT and its activity can be increased by cold exposure. The most common location for BAT in adult humans is the cervical-supraclavicular depot marked by high labeled glucose uptake in that region [Cypess *et al*., 2009; Virtanen *et al*., 2009]. Assessing the morphology of the biopsies’ specimen from supraclavicular depot, adipocytes with numerous multilocular and intracellular lipid droplets were detected. A more recent study improved the PET-computed tomography (CT) method to localize brown/beige adipose depots in humans. Using a refined technique of the PET-CT method, Leitner *et al*. (2017) identified the whole-body BAT distribution and estimated its thermogenic capacity. BAT and brownable adipose depots were found interspersed in several areas, such as cervical, supraclavicular, axillary, mediastinal, paraspinal, and abdominal. Intriguingly, individuals with obesity possessed a higher amount of brownable adipose tissue than lean individuals. However, the amount of active BAT in people with obesity was found to be lower than in lean individuals [Leitner *et al*., 2017].

Research on brown and beige adipocytes has gained an increased attraction in the recent years as they can provide a potential new therapeutic target in the treatment of obesity and various metabolic disorders [Kajimura *et al*., 2015; Sharp *et al*., 2012; Wu *et al*., 2012]. A strong negative correlation was shown between the activity of BAT and the body mass index or the percentage of body fat in humans [van Marken Lichtenbelt *et al*., 2009]. Targeting BAT by enhancing its thermogenic activity or by recruiting new beige adipocytes through the „browning” process can improve glucose utilization and lipid clearance, leading to an improved metabolic balance. With this regard, possible strategies have been raised including central stimulation of the sympathetic innervation of BAT, transplantation of the existing BAT, as well as stem cell therapy [Trayhurn, 2018].

Previously, we reported the global expression profile and thermogenic capacity of adipocytes derived from subcutaneous (SC) and deep neck (DN) biopsies [Toth *et al*., 2020]. Based on the RNA-sequencing analysis, 1049 differentially expressed genes (DEGs) were found in the comparison of SC and DN-derived adipocytes. Among these DEGs, we found 21 solute carrier (SLC) transporters and revealed the importance of alanine-serine-cysteine transporter 1 (ASC1, encoded by *SLC7A10*) [Arianti *et al*., 2021] and thiamine transporter 2 (ThTr2, encoded by *SLC19A3*) [Arianti *et al*., 2022] during thermogenic activation. One of the strongest upregulated SLC transporters in thermogenic DN-derived adipocytes was ThTr2, whose pharmacological inhibition attenuated cAMP-mediated thermogenic activation of human neck area-derived adipocytes [Arianti *et al*., 2022].

Vitamin B1, also known as thiamine, is a water-soluble vitamin that plays a crucial role in carbohydrate, lipid, and protein metabolism [Sriram *et al*., 2012]. Intracellular thiamine is mainly present in its active form, thiamine pyrophosphate (TPP) which is formed by TPP kinase. TPP acts as a coenzyme for several enzymes, such as α-ketoglutarate dehydrogenase or pyruvate dehydrogenase (PDH) therefore contributing to the activity of the tricarboxylic acid cycle. The presence of TPP is also required for transketolase (shifts excess fructose-6-phosphate to glyceraldehyde-3-phosphate) enzyme acting in the pentose phosphate pathway [Lonsdale, 2018]. The level of thiamine in the human body is entirely dependent on dietary intake; there is no known endogenous synthesis, although some types of bacteria in the gut can produce small amounts of thiamine [Teran *et al*., 2021]. The half-life of thiamine is notably short (1-12 hours) and its storage capacity in the body is limited; therefore, continuous dietary supplementation is necessary to maintain tissue thiamine levels at a constant level [Whitfield *et al*., 2018]. The total thiamine concentration in human blood plasma ranges around 10-20 nM, which is remarkably lower as compared to rodents, where it can reach 100 nM [Gangolf *et al*., 2010].

Our current study presents a novel approach to facilitate human adipocyte browning *ex vivo* by providing either gradually increasing or abundant amounts of thiamine during the differentiation of human neck area-derived adipocytes. The presence of gradually increasing concentrations of thiamine in a thiamine-free medium led to elevated basal, cAMP-stimulated, and proton-leak respiration along with increased mitochondrial biogenesis of the differentiated adipocytes. Long-term thiamine treatment also upregulated thermogenesis-related genes and proteins in primary SC and DN-derived adipocytes in an increased proportion to the available thiamine concentration. These results suggest that thiamine availability determines the efficiency of differentiation towards thermogenic cells.

The commercially available culture media contain a much higher thiamine concentration (8.2 μM) than that is found in human blood plasma. Providing excess (25 μM or 50 μM) thiamine further increased the thermogenic competency of SC and DN-derived adipocytes. Our results raise the possibility of a novel strategy of long-term thiamine supplementation, which can elevate the thermogenic competency of differentiating neck area-derived adipocytes for combating obesity.

## 2 Materials and methods

### 2.1 Materials

All chemicals were acquired from Sigma-Aldrich (Munich, Germany) unless otherwise stated.

### 2.2 Ethical statement and obtained human adipose-derived stromal cells (hASCs)

Tissue collection was approved by the Medical Research Council of Hungary (20571-2/2017/EKU) followed by the EU Member States’ Directive 2004/23/EC on presumed consent practice for tissue collection. All experiments were carried out in accordance with the guidelines of the Helsinki Declaration. Written informed consent was obtained from all participants before the surgical procedure. During thyroid surgeries, a pair of DN and SC adipose tissue biopsy samples were obtained to rule out inter-individual variations. Patients with known DM, malignant tumor, or abnormal thyroid hormone levels at the time of surgery were excluded. HASCs were isolated from the stromal-vascular fraction (SVF) of SC and DN fat biopsies as described previously [Tóth *et al*., 2020; Kristóf *et al*., 2019]. The absence of mycoplasma contamination was confirmed by PCR analysis (PCR Mycoplasma Test Kit I/C, Promocell, Heidelberg, Germany).

### 2.3 Differentiation and treatment of hASCs

Human primary adipocytes (ADIP) were differentiated from SVF of adipose tissue containing hASCs (or preadipocytes) according to a described protocol applying cortisol, T3, insulin, dexamethasone and short-term rosiglitazone treatment [Kristóf *et al*., 2015]. We also differentiated preadipocytes under long-term rosiglitazone effect resulting in higher browning capacity of the adipocytes (B-ADIP) [Tóth *et al*., 2020; Elabd *et al*., 2009].

hASCs were differentiated to ADIP or B-ADIP using DMEM-F12-HAM medium with or without the addition of excess (25 μM or 50 μM) thiamine (cat# 731188). In thiamine concentration dependence experiments, we used a custom-made thiamine-free culture fluid (Gibco cat# ME2107367, Waltham, MA, USA) for differentiating the cells supplemented with gradually increasing concentrations (40 nM, 200 nM, 1 μM, 5 μM, or 25 μM) of thiamine. Media were changed every 3 days and cells were lysed after 14 days of differentiation. In every repetition, untreated samples were obtained from the same donor. Cells were incubated at 5% CO_2_ and 37 ^0^C.

### 2.4 RNA isolation and Quantitative Real Time PCR (RT-qPCR)

Cells were collected in TRIzol, total RNA was isolated by chloroform-isopropanol extraction, and RT-qPCR was performed as described previously [Klusóczki *et al*., 2019; Kristóf *et al*., 2016; Arianti *et al*., 2022]. Custom-made gene primers and probes were designed and supplied by Thermo Fisher Scientific (Waltham, MA, USA) as listed in Supplementary Table 1.

### 2.5 Immunoblotting and densitometry

Immunoblotting and densitometry were carried out as described previously [Szatmári-Tóth *et al*., 2020; Arianti *et al*., 2022]. Antibodies and working dilutions are listed in Supplementary Table 2.

### 2.6 Oxygen consumption (OCR) and extracellular acidification rate (ECAR) measurement

OCR and ECAR of adipocytes, which were differentiated as described *in 2*.*3*, were measured using an XF96 oximeter (Seahorse Biosciences, North Billerica, MA, USA) according to previously optimized protocols [Klusóczki *et al*., 2019; Kristóf *et al*., 2016; Arianti *et al*., 2022]. When the adipocytes were differentiated in the presence of excess thiamine, cells were pre-incubated for 1 h and treated in Agilent Seahorse XF Base Medium (Agilent Technologies, cat#103334-100), which is optimized for extracellular flux assays and contains thiamine at 8.2 μM concentration. When the adipocytes were differentiated at thiamine concentrations close to the physiological blood plasma levels, the measurement was performed in thiamine free medium to carry out the assay with the previously obtained thiamine pool of the cells which was gained during differentiation.

In both types of experimental settings, after recording the baseline OCR, 500 μM dibutyril-cAMP was injected to mimic an adrenergic cue; then stimulated OCR was recorded every 30 minutes. Proton-leak respiration was monitored after injecting oligomycin (cat#495455) at 2 μM concentration. Finally, adipocytes received a single bolus of Antimycin A (cat#A8674) at 10 μM concentration for baseline correction (measuring non-mitochondrial respiration). The OCR was normalized to protein content.

### 2.7 Statistical analysis

The results are expressed as mean±SD. The normality of the distribution of the data was tested by Shapiro–Wilk test. For multiple comparisons of each group, one-way ANOVA and Tukey’s post hoc test were used. The data were visualized and analyzed by using GraphPad Prism 8 (GraphPad Software, San Diego, CA, USA).

## 3 Results

### 3.1 Abundant levels of thiamine during adipogenesis elevates the expression of thiamine transporters (ThTrs)

In our previous study, we found that high expression of ThTrs and high thiamine influx contributed to the effective thermogenic activation of neck area-derived adipocytes [Arianti *et al*., 2022]. Therefore, we primarily investigated how the expression of the ThTrs is affected by the presence of excess (25 μM or 50 μM) or gradually increasing (40 nM, 200 nM, 1 μM, or 5 μM) concentrations of thiamine during the differentiation of human SC and DN-derived ADIPs and B-ADIPs. In the first experimental setup, adipocytes were differentiated to ADIPs or B-ADIPs at regular culture conditions; in the presence of 8.2 μM of thiamine found in DMEM-F12-HAM medium, or when excess thiamine was administered to the differentiation regimen. As expected [Arianti *et al*., 2022], ThTr2, but not ThTr1, was expressed at a higher level in DN-derived ADIPs, compared with SC ones, both at mRNA (Fig. 1a-b) and protein levels (Fig. 1c). Focusing on the different applied thiamine concentrations, 25 μM thiamine treatment significantly increased the expression of ThTr1 in SC-derived B-ADIPs, while in DN-derived B-ADIPs, both 25 μM and 50 μM treatments resulted in an increasing effect (Fig. 1a). The excess thiamine did not affect the mRNA expression of ThTr2 (Fig. 1b) and the expression of the investigated ThTr proteins significantly (Fig. 1c).

**Figure 1.**
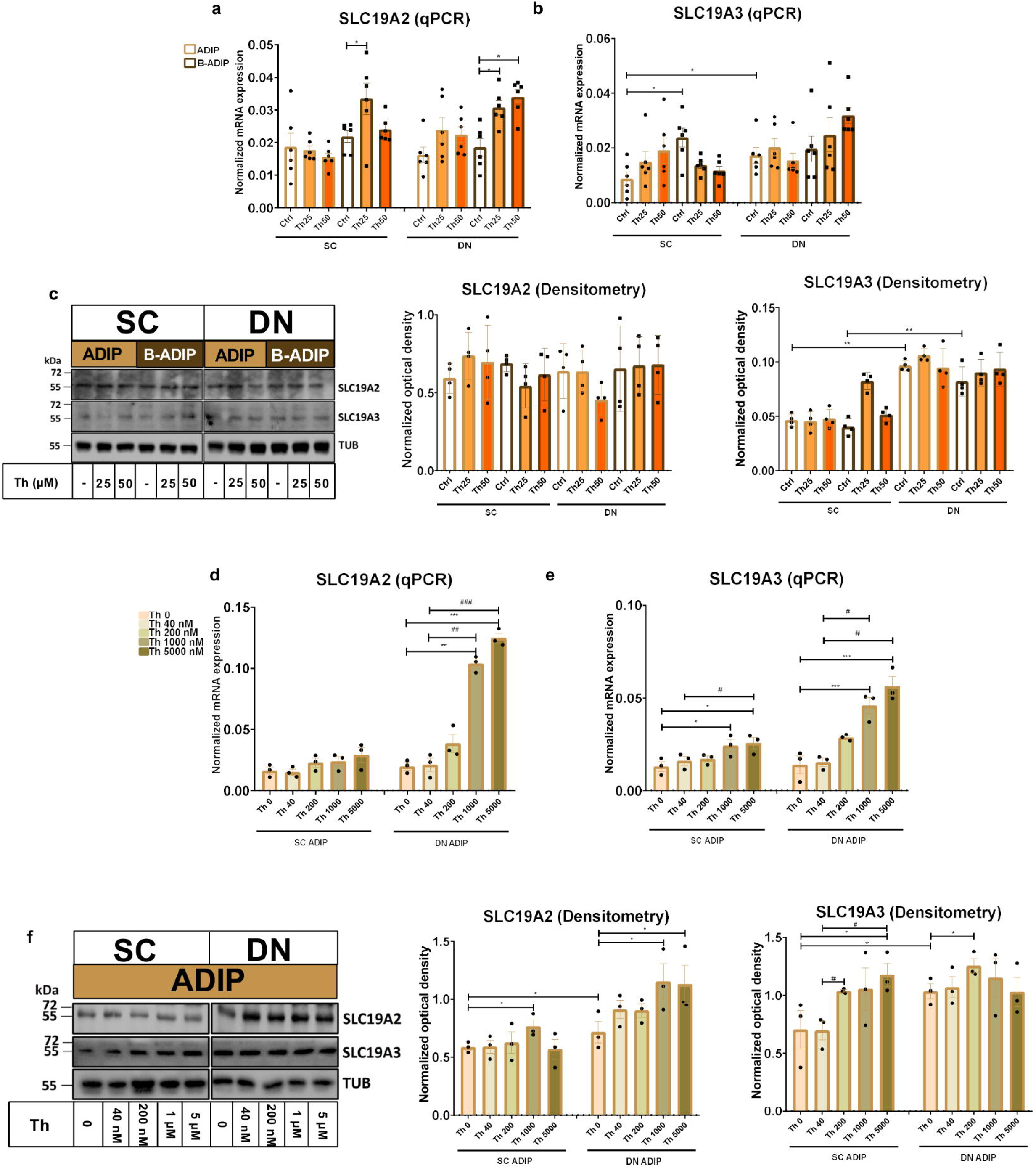
Effect of excess (25 μM and 50 μM) and gradually increasing (40 nM, 200 nM, 1 μM, 5 μM) concentrations of thiamine (Th) on the expression of Th transporters (ThTrs) in human subcutaneous (SC) and deep neck (DN)-derived adipocytes. ADIPs and B-ADIPs were differentiated at regular culture conditions (Ctrl, 8.2 μM Th) or in the presence of 25 μM (Th25) or 50 μM (Th50) of Th (a-c). (a-b) mRNA expression of *SLC19A2* and *SLC19A3* assessed by RT-qPCR, n=6. (c) ThTr1 (SLC19A2) and ThTr2 (SLC19A3) protein expression detected by immunoblotting, n=4. SC or DN-derived preadipocytes were differentiated into ADIPs in Th-free medium (Th 0) or in the presence of gradually increasing concentrations of Th (40 nM, 200 nM, 1 μM, 5 μM) (d-f). (d-e) mRNA expression of *SLC19A2* and *SLC19A3* assessed by RT-qPCR, n=3. (f) ThTr1 (SLC19A2) and ThTr2 (SLC19A3) protein expression detected by immunoblotting, n=3. Statistical analysis was performed by one-way ANOVA, *#p<0.05, **##p<0.01, ***###p<0.001, *comparing data at each concentration of Th to the lack of Th or # comparing the indicated groups.

In the second experimental setup, we studied the importance of thiamine availability even at ranges corresponding to physiological levels during adipocyte differentiation, where SC or DN-derived hASCs were differentiated to ADIPs or B-ADIPs in thiamine-free medium (Th 0) or when thiamine was re-administered at gradually increasing concentrations. The mRNA expression of both ThTrs was not affected by the anatomical origin of the ASCs when ADIPs were differentiated in the presence of 200 nM or less thiamine, however, it was upregulated in direct proportion to increasing thiamine concentrations, where the presence of 1 μM or 5 μM thiamine resulted in a significant effect compared to Th 0 condition or to the 40 nM concentration, which is close to the physiological concentration of the vitamin found in human blood plasma (Fig. 1d-e). A similar trend was observed when the protein expression of the ThTrs was investigated (Fig. 1f). In the case of B-ADIPs, the anatomical origin of the progenitors did not affect the expression of the ThTrs at Th 0 condition, however, both ThTrs mRNA were expressed at a higher level in DN-derived as compared to SC ones when B-ADIPs were differentiated in the presence of 40 nM thiamine. Similarly to our observations in ADIPs, thiamine potentiated the mRNA expression of both ThTrs in B-ADIPs, especially in DN-derived ones, in a concentration-dependent manner (Fig. S1a). In summary, extracellular thiamine levels during the differentiation of neck area-derived adipocytes positively associate with the expression of ThTrs, and adipocytes differentiated at regular culture conditions are fully equipped with ThTrs. The expression of *SLC25A19* that encodes the mitochondrial TPP transporter was not significantly affected by the anatomical origin of the hASCs or by the availability of thiamine at physiological (Fig. S1b) or at abundant levels (Fig. S2b) in the differentiation media of adipocytes.

### 3.2 Thiamine availability determines the thermogenesis of differentiating adipocytes in a concentration-dependent manner

Next, we studied the importance of thiamine availability in the thermogenesis of ADIPs which were differentiated either in Th 0 medium or in the presence of gradually increasing concentrations of thiamine (40 nM, 200 nM, 1 μM, 5 μM, or 25 μM) re-administered to the Th 0 medium. After recording basal OCR, dibutyryl-cAMP, a cell-permeable cAMP analog that imitates adrenergic stimulation of thermogenesis, was injected, which strongly increased mitochondrial respiration, as reported previously [Tóth *et al*., 2020]. Then proton-leak respiration, an indicator of UCP1-dependent thermogenesis, was measured by the administration of oligomycin, which inhibits ATP synthase. Basal OCR, the maximal respiration rate after the injection of the cAMP analog, and stimulated proton-leak OCR tended to elevate in proportion to the increasing thiamine concentrations within the differentiation media from 40 nM up to 25 μM in both SC and DN-derived ADIPs compared to the Th 0 condition. In the case of SC-derived ADIPs, statistically significant differences were observed from 200 nM thiamine availability (Fig. 2a), while in DN-derived ADIPs, only the differences between 25 μM and 40 nM thiamine treatments or Th 0 controls were statistically significant, respectively (Fig. 2b). Of note, the extracellular flux assay was carried out in Th 0 medium to measure OCR with the previously obtained thiamine pool of the adipocytes which was gained during differentiation.

**Figure 2.**
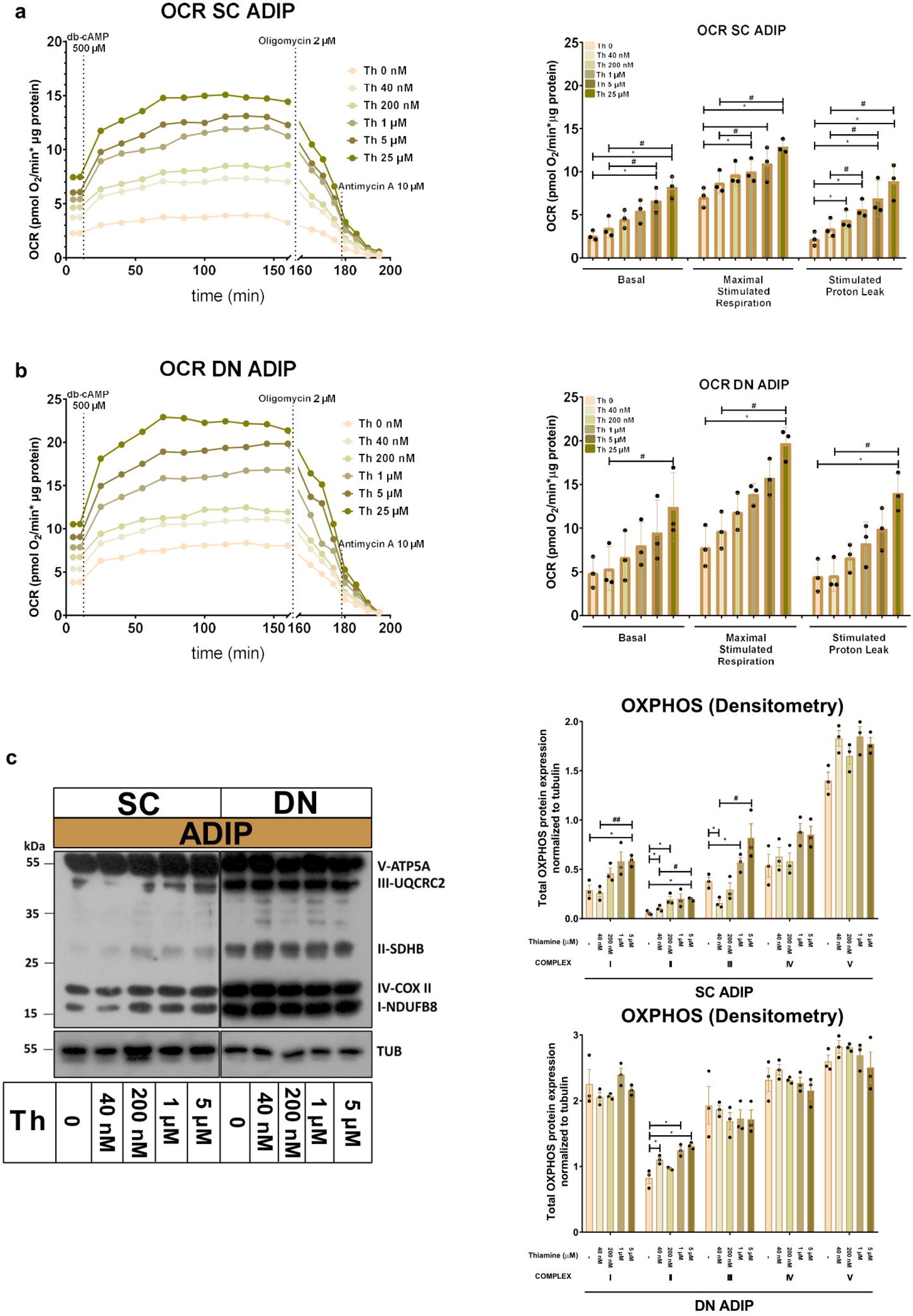
Gradually increasing thiamine (Th) concentrations during differentiation increases oxygen consumption rate (OCR) and the expression of OXPHOS complex subunits of human subcutaneous (SC) and deep neck (DN)-derived adipocytes (ADIPs). (a) SC or (b) DN-derived preadipocytes were differentiated into ADIPs in Th-free medium (Th 0) or in the presence of gradually increasing concentrations of Th (40 nM, 200 nM, 1 μM, 5 μM, 25 μM), then OCR was detected at basal conditions, following the injection of 500 μM dibutyryl-cAMP for 6 hours, and after 2 μM oligomycin administration (stimulated proton-leak respiration). Each left panel shows representative curves of 3 measurements with independent donor samples. OCR at basal, maximal stimulation post cAMP injection, and after oligomycin administration (right panels) were quantified in ADIPs derived from three independent donors. (c) Total OXPHOS protein subunit expression detected by immunoblotting, n=3. Statistical analysis was performed by one-way ANOVA, *#p<0.05, **#p<0.01, *comparing data at each concentration of thiamine to the lack of Th (Th 0) or # comparing the indicated groups.

The protein expression of mitochondrial complex subunits I, II, and III showed an increasing tendency in SC-derived ADIPs differentiated in the presence of gradually increasing concentrations of thiamine, compared to the Th 0 control. A statistically significant increase was observed in the case of 1 μM and 5 μM thiamine administration compared to 40 nM thiamine treatment or Th 0, respectively. The amount of complex subunits IV and V was not affected by thiamine availability up to 5 μM in SC-derived ADIPs. Only the expression of complex subunit II was influenced by the extracellular thiamine levels in DN-derived ADIPs, which already expressed these proteins at large amounts (Fig. 2c). These results suggest that thiamine availability during adipocyte differentiation has a significant effect on the thermogenic competency of neck-derived ADIPs.

### 3.3 Increasing concentrations of thiamine during adipocyte differentiation enhances thermogenic gene expression in adipocytes

As we observed that increasing concentrations of thiamine in the adipogenic differentiation media positively associated with the mitochondrial respiration, especially with proton-leak OCR, of neck area-derived ADIPs, we also investigated how the availability of thiamine affects the expression of thermogenesis-related genes. *UCP1* and *PGC1A* mRNA expression increased in direct proportion to the increasing thiamine concentrations in both SC and DN-derived ADIPs (Fig. 3a-b), statistically significant effects were observed from 40 nM or 1 μM thiamine availability as compared to Th 0, respectively. In the case of B-ADIPs, the same trend was found, with statistically significant changes from 200 nM thiamine administration as compared to Th 0 with regard to both genes (Fig. S2a-b). In SC-derived ADIPs, UCP1 but not PGC1A protein expression was significantly affected by thiamine availability in a concentration-dependent manner (Fig. 3c, Fig. S3a). In SC-derived B-ADIPs, DN-derived ADIPs, and DN-derived B-ADIPs, the expression of both investigated proteins was increased when the thiamine levels were consecutively raised during adipogenesis. In these cases, the effect of thiamine availability was statistically significant even at concentrations corresponding to physiological human blood plasma thiamine levels (Fig. 3c, Fig. S2c, Fig. S3a).

**Figure 3.**
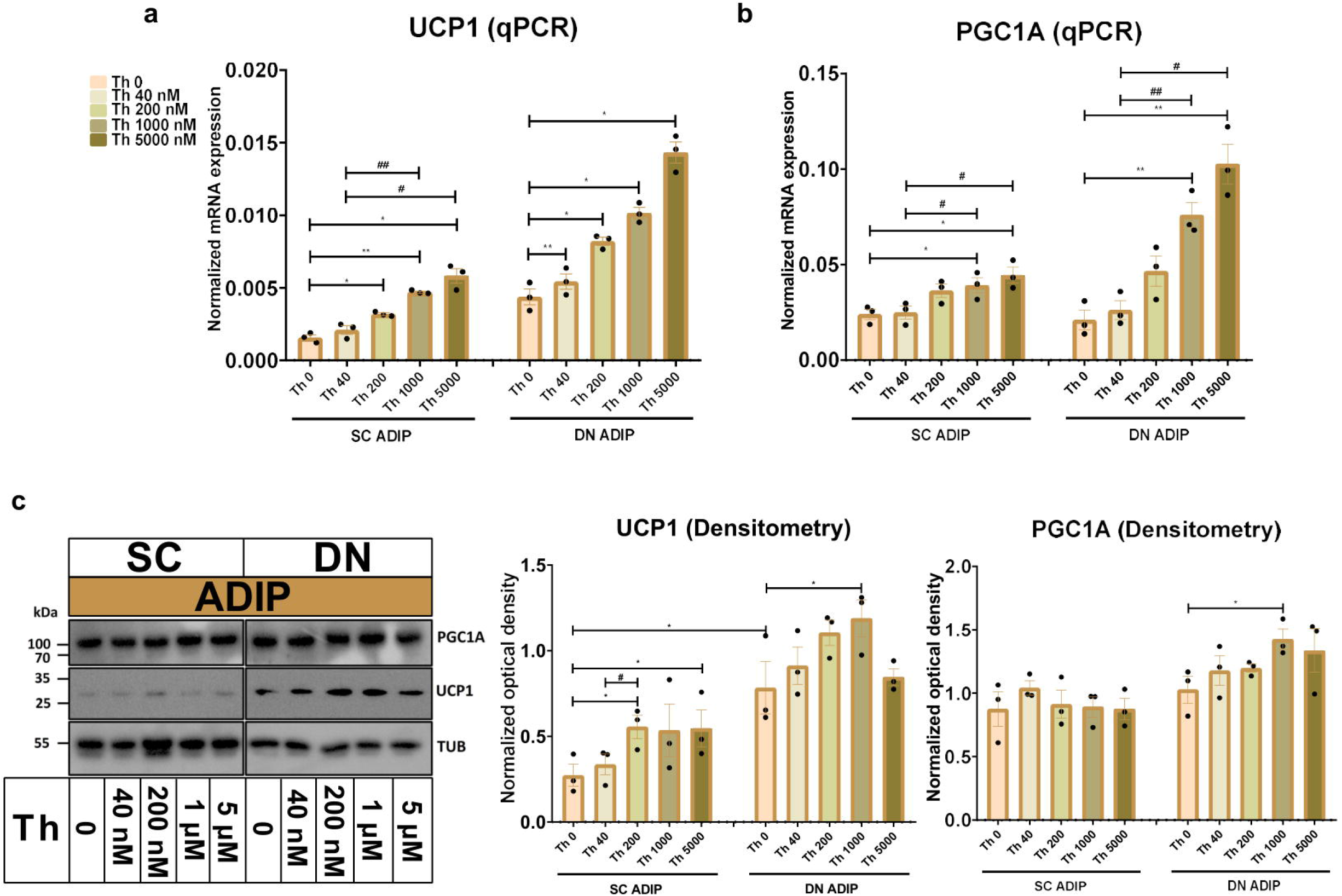
Effect of gradually increasing concentrations of thiamine (Th) on thermogenic gene and protein expression in human subcutaneous (SC) and deep neck (DN)-derived adipocytes. ADIPs were differentiated and treated as in Figure 2. (a-b) mRNA expression of *UCP1* and *PGC1A* assessed by RT-qPCR, n=3. (c) UCP1 and PGC1A protein expression detected by immunoblotting, n=3. Uncropped images are shown in Figure S3a. Statistical analysis was performed by one-way ANOVA, *#p<0.05, **#p<0.01, *comparing data at each concentration of Th to the lack of Th (Th 0) or # comparing the indicated groups.

We also investigated the effect of thiamine availability on the mRNA expression of several thermogenesis-related genes and the white marker *LEP* in SC and DN-derived ADIPs (Fig. 4) and B-ADIPs (Fig. S4). Increased amounts of extracellular thiamine upregulated the expression of *DIO2* [Canon and Nedergaard, 2004], *TBX1* [Wu *et al*., 2012], *CKMT2* [Kazak *et al*., 2015], and *CIDEA* [Garcia *et al*., 2016] in a concentration-dependent manner in ADIPs and B-ADIPs of both origins. *CITED1* [Sharp *et al*., 2012] was expressed at the highest level when both types of adipocytes were differentiated in the presence of 40 nM thiamine. In SC and DN-derived ADIPs, Th 0 condition and higher availability of thiamine tended to decrease *CITED1* mRNA expression, while in the case of B-ADIPs, the beige marker gene was expressed significantly lower in these conditions. *LEP* mRNA expression showed a similar trend, however, we observed suppressive effects at statistically significant levels of higher doses of thiamine in DN ADIPs as well. Altogether, our data suggest that thiamine availability during adipocyte differentiation, even at concentrations found in human blood plasma, has a significant impact on the expression of thermogenesis-related genes.

**Figure 4.**
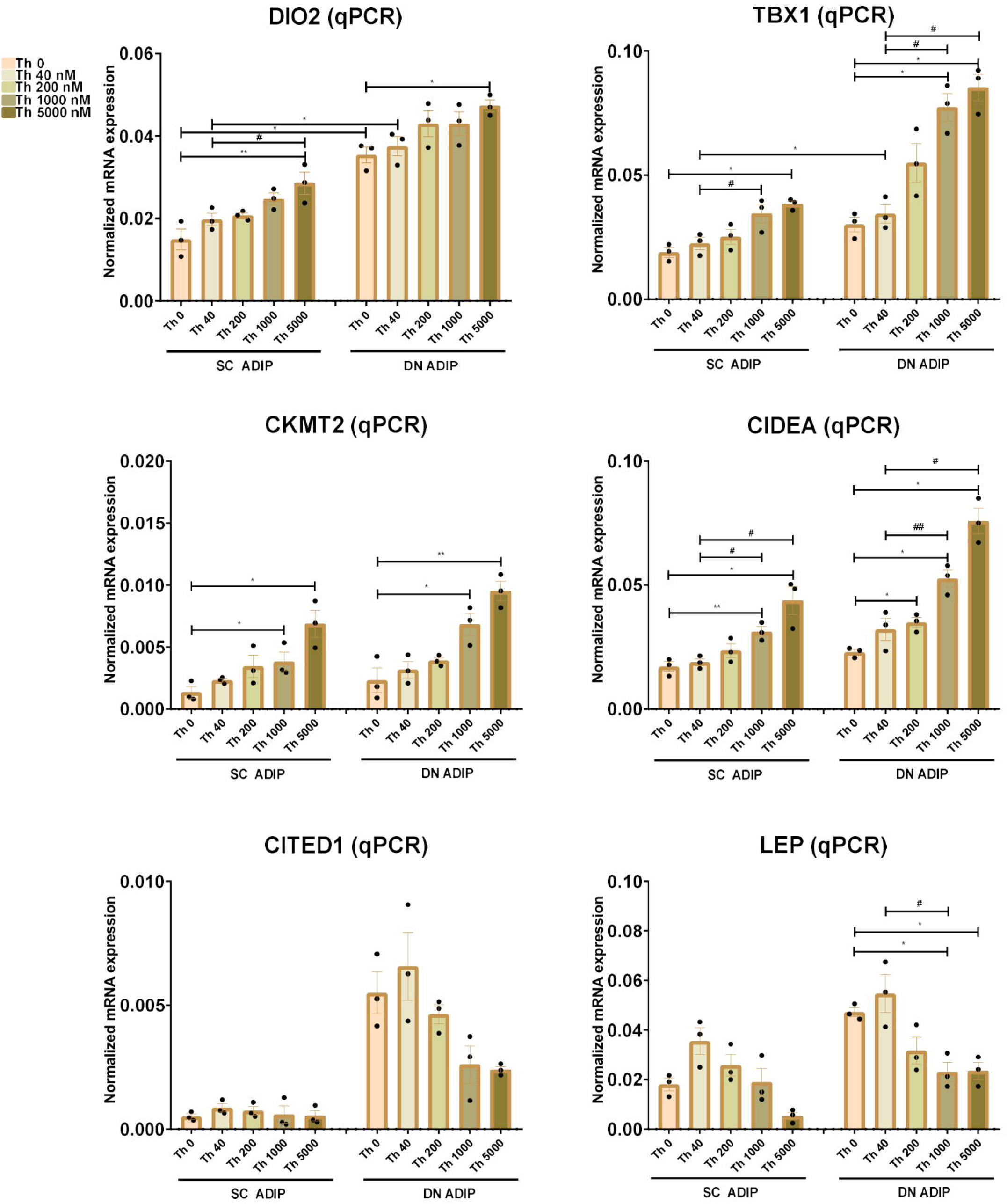
Effect of gradually increasing concentrations of thiamine (Th) on thermogenic gene induction in human subcutaneous (SC) and deep neck (DN)-derived adipocytes. ADIPs were differentiated and treated as in Figures 2 and 3. (a-f) mRNA expression of *DIO2, TBX1, CKMT2, CIDEA, CITED1*, and *LEP* assessed by RT-qPCR, n=3. Statistical analysis was performed by one-way ANOVA, *#p<0.05, **##p<0.01, *comparing data at each concentration of Th to the lack of Th (Th 0) or # comparing the indicated groups.

### 3.4 Excess thiamine treatment during adipocyte differentiation enhances uncoupled respiration

Next, we investigated how the administration of abundant amounts of thiamine to the regular adipocyte differentiation medium (that already contained 8.2 μM of thiamine) affected the thermogenesis of SC and DN-derived ADIPs. SC-derived ADIPs differentiated in the presence of 25 μM excess thiamine had significantly higher basal OCR compared to those differentiated with regular differentiation medium (Fig. 5a, left and middle panels). In the case of DN-derived ADIPs, a similar trend was observed without statistical significance (Fig. 5b, left and middle panels). The presence of excess thiamine in the adipogenic differentiation cocktail resulted in higher maximal cAMP-stimulated respiration in ADIPs of both SC and DN origins. Both SC and DN-derived unstimulated ADIPs had higher proton-leak OCR when the hASCs were differentiated in the presence of excess thiamine. As expected, thermogenic stimulation with dibutyryl-cAMP elevated proton-leak respiration, especially in DN-derived ADIPs. In the case of stimulated ADIPs, the difference caused by excess thiamine availability during differentiation was observed only in SC but not in DN-derived cells at a statistically significant level (Fig. 5a-b, left and middle panels). The ECARs were also elevated in response to stimulation in both SC and DN-derived ADIPs, however, were not affected by the thiamine concentrations applied during adipogenesis (Fig. 5a-b, right panels).

**Figure 5.**
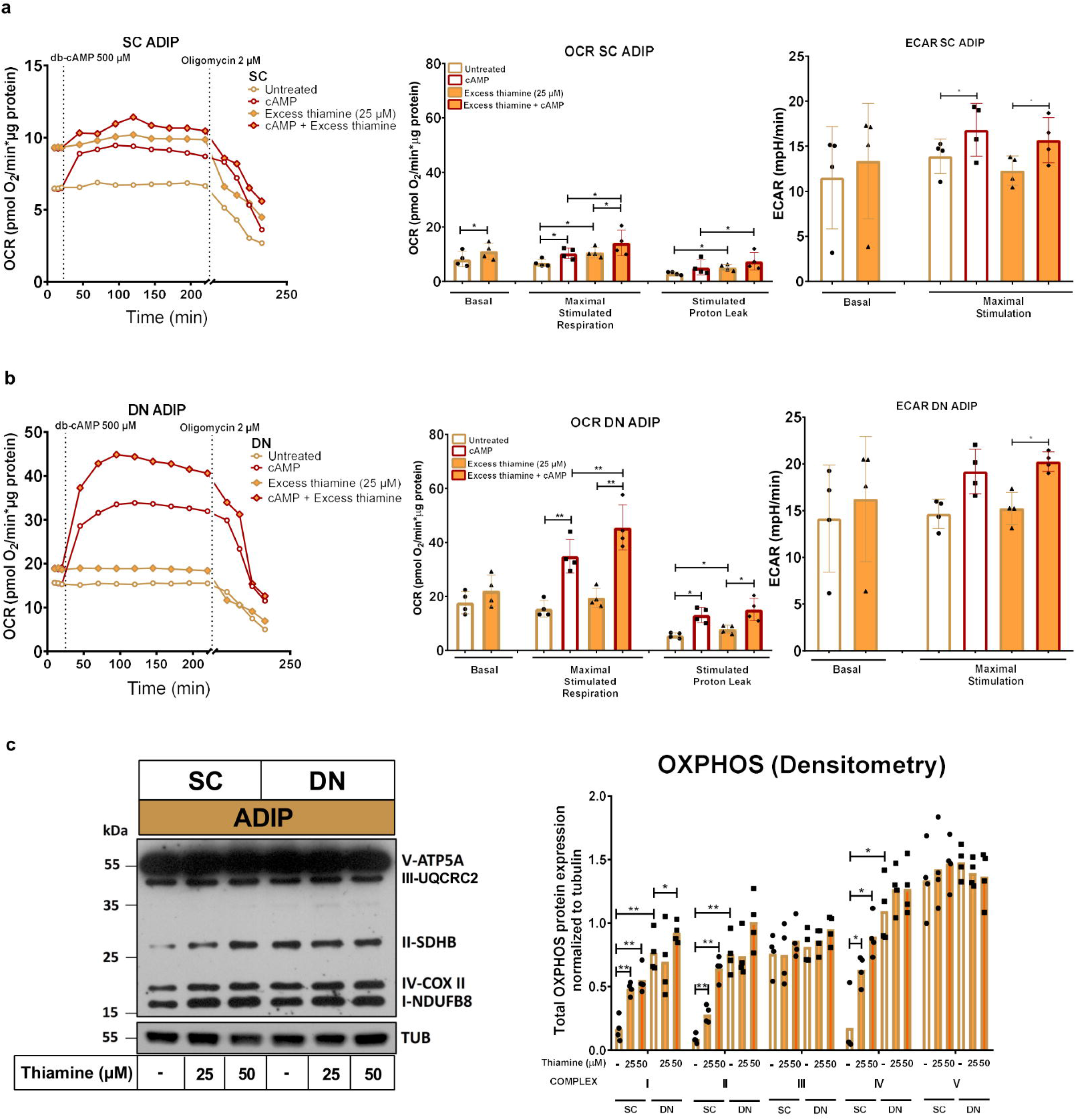
Excess thiamine provided at adipogenic differentiation increases oxygen consumption rate (OCR) and the expression of OXPHOS complex subunits of human subcutaneous (SC) and deep neck (DN)-derived adipocytes (ADIPs). (a) SC or (b) DN-derived preadipocytes were differentiated into ADIPs at regular culture conditions (8.2 μM thiamine, white bars) and in the presence of excess thiamine (25 μM, orange bars), then OCR was detected at basal conditions, following the injection of 500 μM dibutyryl-cAMP for 6 hours, and after 2 μM oligomycin administration (proton-leak respiration). Each left panel shows representative curves of 4 measurements with independent donor samples. OCR at basal, maximal stimulation post cAMP injection, and after oligomycin administration (middle panels), and extracellular acidification rate (ECAR) (right panels) were quantified in ADIPs derived from 4 independent donors. (c) Total OXPHOS protein subunit expression detected by immunoblotting, n=4. Statistical analysis was performed by one-way ANOVA, *p<0.05, **p<0.01.

The protein expression of mitochondrial complex subunits I, II, and IV were significantly increased upon 25 μM or 50 μM of thiamine treatment in SC-derived differentiating ADIPs, while 50 μM thiamine administration resulted in higher complex I and II subunit amounts in DN-derived ADIPs which had generally higher complex subunits I, II, and IV expression as compared to the SC ones (Fig. 5c). These results demonstrate that excess thiamine present in large amounts during adipocyte differentiation further increases the thermogenic competency and mitochondrial biogenesis of neck area-derived ADIPs.

### 3.5 Abundant thiamine during adipocyte differentiation elevates thermogenic gene expression

After observing an elevation in the proton-leak respiration in the presence of excess thiamine, we hypothesized that abundant thiamine during adipocyte differentiation can further elevate the expression of thermogenic and mitochondrial biogenesis-related genes. We found that at mRNA levels, in the case of both *UCP1* (Fig. 6a) and *PGC1A* (Fig. 6b), the presence of excess thiamine increased their expression compared to the control ADIPs or B-ADIPs. In accordance with previous observations [Tóth *et al*., 2020], the expression of these markers was higher in DN-derived and B-ADIPs compared to SC-derived or ADIPs, respectively. 50 μM of thiamine resulted in stronger effects in several cases in both ADIPs and B-ADIPs derived from both anatomical origins (Fig. 6a-b). At protein levels, we observed that UCP1 expression was increased in the presence of excess thiamine (both 25 μM and 50 μM) compared to the controls, which was in association with the mRNA expression. These differences were statistically significant in the case of B-ADIPs. With regard to PGC1A, we observed a significant difference caused by the PPARγ-driven B-ADIP differentiation in SC-derived cells, however, abundant thiamine administration did not further increase the protein expression of the mitochondrial biogenesis master regulator (Fig. 6c, Fig. S3b).

**Figure 6.**
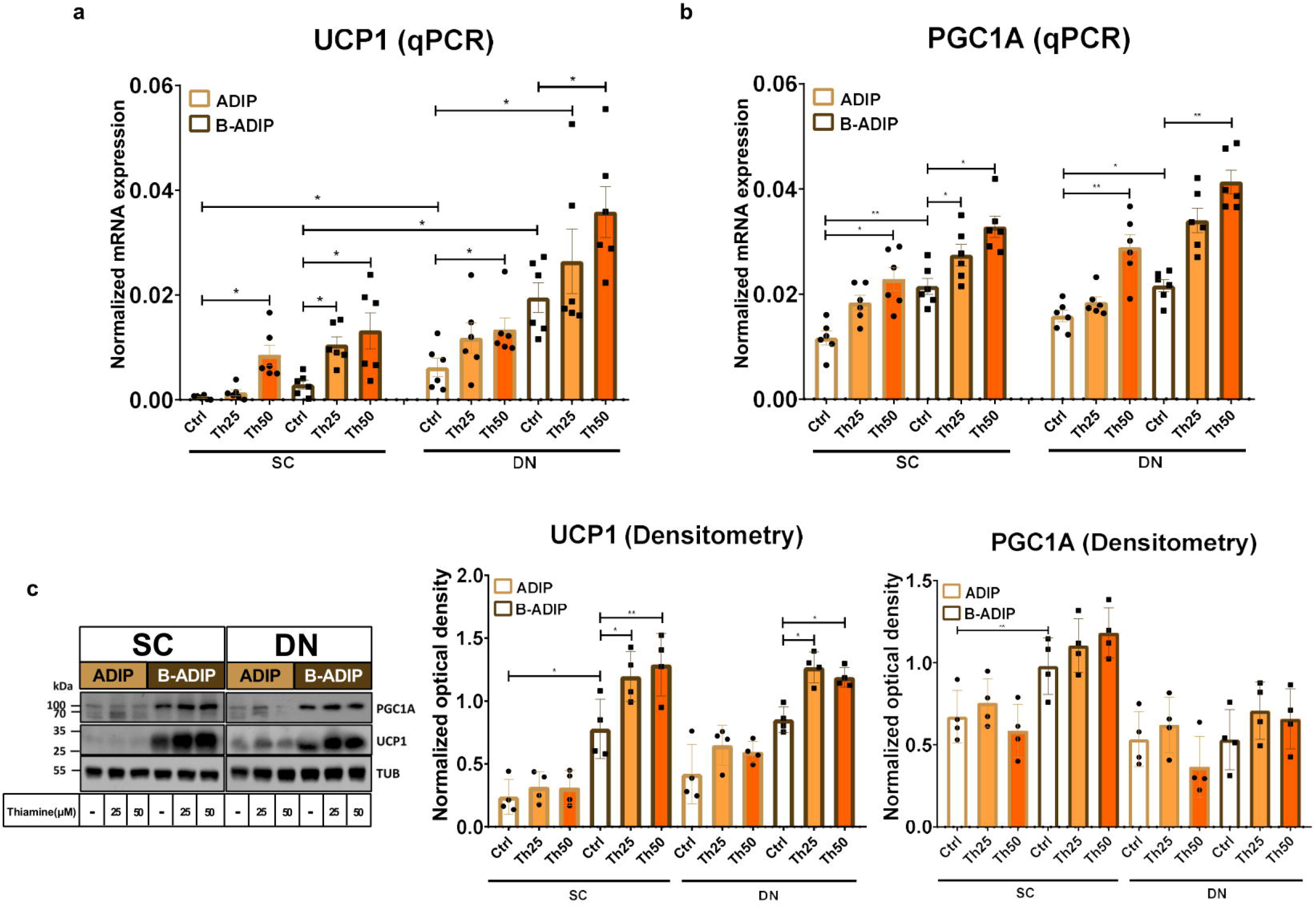
Excess thiamine (Th) during adipogenic differentiation increases the expression of thermogenesis markers in human subcutaneous (SC) and deep neck (DN)-derived adipocytes. ADIPs and B-ADIPs were differentiated at regular culture conditions (Ctrl, 8.2 μM Th, white bars) or in the presence of 25 μM (Th25) or 50 μM (Th50) of excess Th (orange bars). (a-b) mRNA expression of *UCP1* and *PGC1A* assessed by RT-qPCR, n=6. (c) UCP1 and PGC1A protein expression detected by immunoblotting, n=4. Uncropped images are shown in Figure S3b. Statistical analysis was performed by one-way ANOVA, *p<0.05, **p<0.01.

We also investigated the effect of excess thiamine administered to the differentiation cocktail on the mRNA levels of *CKMT2, CIDEA, DIO2, TBX1, CITED1*, and *LEP* and found that – with the exception of the white adipogenic marker, *LEP* - thiamine potentiated the expression of these genes in both SC and DN-derived ADIPs and B-ADIPs (Fig. 7). These results suggest that excess thiamine provided during differentiation of neck area-derived adipocytes potentiates the expression of thermogenesis-related and brown/beige marker genes.

**Figure 7.**
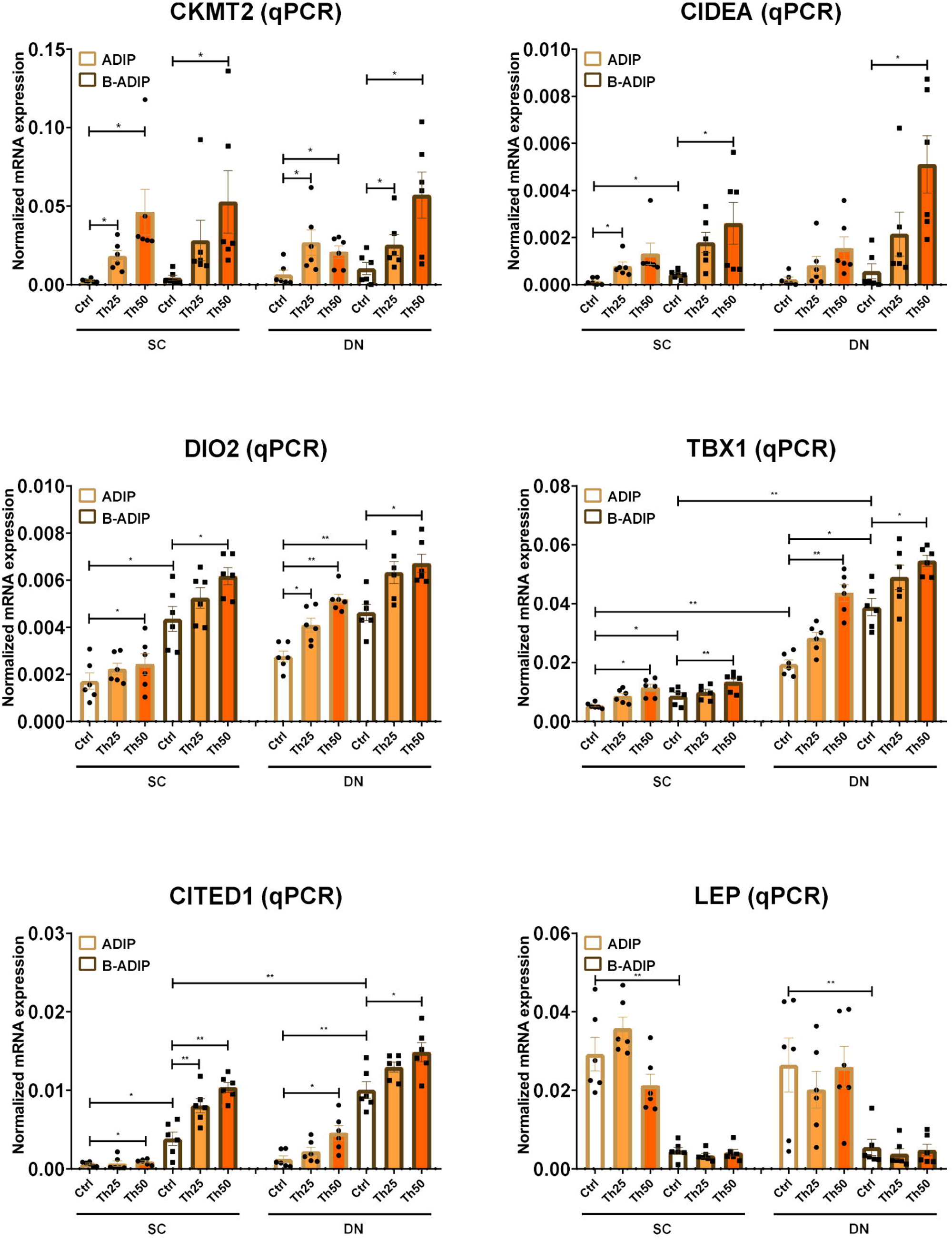
Excess thiamine (Th) during adipogenic differentiation elevates the expression of thermogenesis marker genes in human subcutaneous (SC) and deep neck (DN)-derived adipocytes. ADIPs and B-ADIPs were differentiated and treated as in Figure 6. mRNA expression of *CKMT2, CIDEA, DIO2, TBX1, CITED1*, and *LEP* was assessed by RT-qPCR, n=6. Statistical analysis was performed by one-way ANOVA, *p<0.05, **p<0.01.

## 4 Discussion

To understand the different molecular mechanisms involved in the regulation of thermogenesis, Tóth *et al*. (2020) previously studied differentiated adipocyte populations derived from the SVFs of paired DN and SC adipose tissue sections of nine independent donors and compared their global gene expression patterns using RNA-sequencing. Adipocytes were differentiated by using a standard adipogenic differentiation medium (ADIP) or long-term rosiglitazone treatment (B-ADIP), which enhanced the rate of adipocyte browning. Brown adipocyte content and browning capacity, which were estimated by BATLAS [Perdikari *et al*., 2018] and ProFAT [Cheng *et al*., 2018] open access webtools, respectively, were higher in DN as compared to SC-derived adipocytes irrespective of the applied differentiation protocols [Tóth *et al*., 2020]. Among these DEGs, we found 21 SLC transporters, which mediate nutrient uptake and are known to play role in various biological processes. We have reported that pharmacological inhibition of ThTrs attenuated the efficiency of thermogenic activation in human neck area-derived adipocytes. On the other hand, abundant thiamine levels were required for the effective adrenergic-driven thermogenic stimulation of those cells [Arianti *et al*., 2022]. In this study, we addressed the question how extracellular thiamine availability during differentiation affects the thermogenic capacity of human neck area-derived adipocytes.

Certain types of meals are rich in thiamine including brown rice, whole grains, legumes, and meats [Sharma *et al*., 2013]. The recommended daily allowance (RDA) of thiamine for adults is 1.2 mg/day for males and 1.1 mg/day for females (RDA increases for pregnant women) [Turck *et al*., 2016]. Although thiamine is abundant in several food ingredients commonly consumed diets can be low in micronutrients. Thiamine deficiency, which mostly affects the nervous system, heart, and gastrointestinal tract, usually occurs when the RDA is not maintained. It can be developed because of impaired intestinal absorption or elevated excretion rates of thiamine caused by alcohol dependency or malnutrition [Fouarge and Maquet, 2019; Thomson *et al*., 1970]. The most common diseases resulting from thiamine deficiency are the Beriberi disorder and the Wernicke-Korsakoff syndrome (WKS) [Mateos-Diaz *et al*., 2022]. The main symptoms of WKS are memory impairment, ataxia, and hypothermia, which is frequently reported as a secondary symptom. It has been reported that thiamine deficiency leads to the lesion of the hypothalamus, which is the main regulatory site of body temperature and appetite [Tanev *et al*., 2008]. Human case studies have reported that the hypothermic condition can be treated by thiamine administration for 2 days [Hansen *et al*., 1984]. Our presented data showed that the absence of thiamine during adipogenic differentiation led to a reduced thermogenic capacity of human neck area-derived adipocytes marked by lower cellular respiration, proton-leak respiration reflecting UCP1-dependent heat production, and expression of thermogenic markers. Gradually increasing concentrations of thiamine during differentiation elevated the thermogenic capacity and expression of thermogenic markers in a concentration-dependent manner suggesting the beneficial effect of thiamine in directly boosting the browning capacity of human neck area-derived adipocytes.

Thiamine deficiency is rarely found in developed countries, however, it has become common among people with alcoholism in Europe, North America, and Australia [DiNicolantonio *et al*., 2018]. Thiamine deficiency was also found in 15.5-29% of patients with obesity prior to bariatric surgery [Kerns *et al*., 2015, Carrodeguas *et al*., 2005]. The complications of thiamine deficiency are also found in patients with DM contributing to hyperglycemia-induced cellular damage, endothelial dysfunction, and elevated oxidative stress [DiNicolantonio *et al*., 2018]. Reduced activity of erythrocyte transketolase was found in 17-79% of patients with DM [Jermendy *et al*., 2006; Page *et al*., 2011]. Another study involving 120 adults with type 2 DM (out of which 46 individuals had microalbuminuria) reported that thiamine deficiency occurred in 98 patients and 100% of total involved subjects with or without microalbuminuria, respectively [Nix *et al*., 2015]. Significantly lower levels of thiamine were also found in 76% or 75% of patients with type 1 or type 2 DM, respectively [Thornalley *et al*., 2007]. Thiamine supplementation (100 mg, 3×100 mg daily which is approximately 100x higher than the RDA) for 6 weeks improved glucose tolerance in individuals with hyperglycemia [Alaei *et al*., 2013]. In drug-naïve patients with type 2 DM, a daily dose of 150 mg of thiamine for one month led to a significant decrease in plasma fasting glucose concentration [Gonzalez-Ortiz *et al*., 2011]. It has also been reported that thiamine supplementation may provide beneficial effects in patients with type 2 DM by improving lipid and creatinine profiles [Al-Attas *et al*., 2014]. Other risk factors of thiamine deficiency are immune diseases, hyperthyroidism, heart failure, and renal failure requiring dialysis [DiNicolantonio *et al*., 2018].

Thiamine, which is converted to TPP, plays an important role in the proper function of mitochondrial enzyme complexes, including PDH. Tanaka *et al*. (2010) reported that male Otsuka Long-Evans Tokushima-Fatty (OLETF) rats treated with water containing 0.2% of thiamine had higher hepatic PDH activity and decreased lipid oxidation. In addition, thiamine treatment prevented obesity and metabolic disorders in OLETF rats by decreasing the body weight and visceral WAT mass and alleviated hypertrophy of epididymal white adipocytes [Tanaka *et al*., 2010].

Hitherto, the effects of micronutrient availability on the differentiation of human adipocytes remain elusive to date. Hirata *et al*. (2020) reported that thiamine promotes the differentiation of T lymphocytes via TGF-β superfamily production in thymic stromal cells suggesting the role of thiamine as a key nutritional regulator for immune cell development. A randomized controlled trial involving 100 individuals with obesity or overweight reported that a high protein diet combined with high thiamine administration (2.8 mg/day) led to reduced body weight up to 11 kg after 16 weeks and without affecting erythrocyte thiamine levels [Keogh *et al*., 2012].

The concentration of thiamine in the regular cell culture medium is close to 10 μM, whereas thiamine concentration in human blood plasma is in the nM range. Our current data revealed that 25 μM thiamine addition to regular cell culture medium during the differentiation of the same adipocyte types elevated basal, maximal, and proton-leak respiration that reflects heat production. Our results also indicated that thiamine supplementation during adipogenesis augmented the expression of thermogenic markers in the thermogenically prone neck area-derived adipocytes *ex vivo*. Brown/beige adipocytes have been proposed as a potential target to treat obesity and DM, mainly due to their high capability as glucose and fatty acid sink [Cypess *et al*., 2009; Wu *et al*., 2006]. Transplantation of hASCs, which are differentiated into brown/beige adipocytes, into recipient mice exerted beneficial effects, such as significant glucose consumption and increased energy expenditure [Singh *et al*., 2020]. Despite the successful differentiation of hASCs into beige adipocytes, increasing the thermogenic capacity of these beige cells to be sufficient to drive heat production *in vivo* remains challenging. Our presented results showed that the addition of extracellular thiamine during the differentiation stage increased proton-leak respiration reflecting high heat production even at basal conditions, which may provide an approach to elevate the thermogenesis of beige adipocytes before transplanting them into the recipient. Therefore, our results highlight that extracellular thiamine availability should be optimized when thermogenic adipocytes are differentiated for transplantation approaches. On the other hand, constant supplementation with thiamine might open up a possibility for a safe and cost-effective nutritional approach for the prevention or treatment of obesity and related metabolic disorders.

## 6 Author Contributions

BÁV and RA performed the experiments. EK, RA, and BÁV conceptualized the research, with inputs from LF. FG provided adipose tissue samples. EK and LF supervised the research and acquired funding. BÁV and RA wrote the original draft of the manuscript. EK and LF edited the final version of the manuscript.

## 8 Acknowledgments

We thank Dr. Éva Csősz for her exceptional help in reviewing the manuscript before its submission and Jennifer Nagy for technical assistance.

## 10 Data Availability Statement

The original contributions presented in this study are included in the article/Supplementary material, further inquiries can be directed to the corresponding author.

