## Supplementary material for "Extracellular thiamine concentration influences thermogenic competency of differentiating neck area-derived human adipocytes"

### 1 Supplementary Figures and Tables

#### 1.1 Supplementary Figures

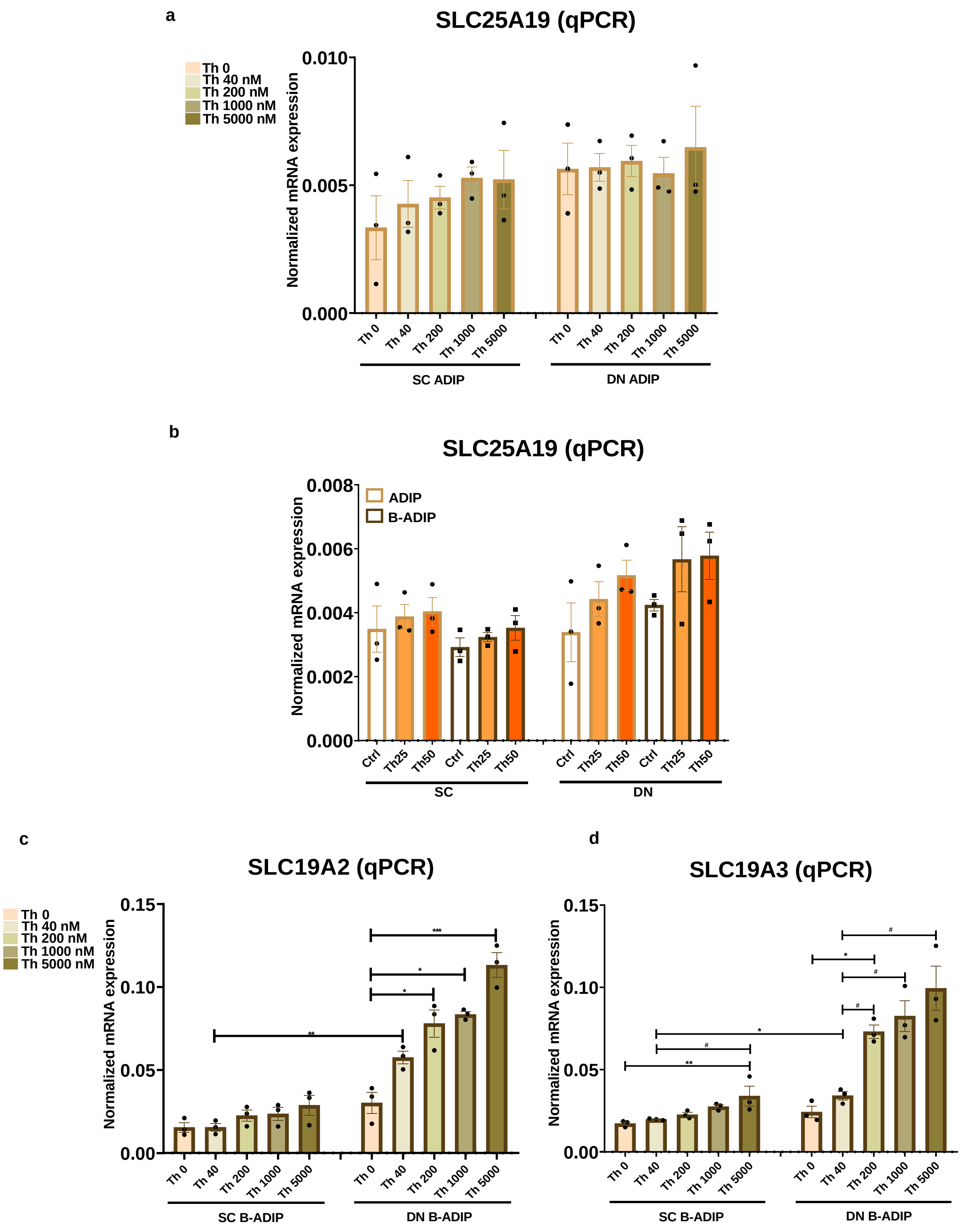
a
SLC25A19 (qPCR)
Th 0
Th 40 nM
Th 200 nM
Th 1000 nM
Th 5000 nM
Normalized mRNA expression
0.010
0.005
0.000
Th 0
Th 40
Th 200
Th 1000
Th 5000
SC ADIP
DN ADIP
b
SLC25A19 (qPCR)
ADIP
B-ADIP
Normalized mRNA expression
0.008
0.006
0.004
0.002
0.000
Ctrl
Th25
Th50
SC
DN
c
SLC19A2 (qPCR)
Th 0
Th 40 nM
Th 200 nM
Th 1000 nM
Th 5000 nM
Normalized mRNA expression
0.15
0.10
0.05
0.00
***
*
*
**
SC B-ADIP
DN B-ADIP
d
SLC19A3 (qPCR)
Normalized mRNA expression
0.15
0.10
0.05
0.00
#
*
#
#
*
#
**
SC B-ADIP
DN B-ADIP

**Supplementary Figure 1.**

Effect of gradually increasing concentrations of thiamine (Th) on the expression of Th transporters in human subcutaneous (SC) and deep neck (DN)-derived brown differentiated adipocytes (B-ADIPs). (a) mRNA expression of *SLC19A2* and *SLC19A3* assessed by RT-qPCR, n=3.

Effect of gradually increasing (40 nM, 200 nM, 1 μM, 5 μM) and excess (25 μM and 50 μM) concentrations of Th on the expression of mitochondrial Th pyrophosphate transporter (encoded by *SLC25A19*) in human SC and DN-derived adipocytes (ADIPs) and B-ADIPs. (c-d) mRNA expression of *SLC25A19* assessed by RT-qPCR, n=3. Statistical analysis was performed by one-way ANOVA, *#p<0.05, **##p<0.01, ***###p<0.001, *comparing data at each concentration of Th to the lack of Th (Th 0) or # comparing the indicated groups.

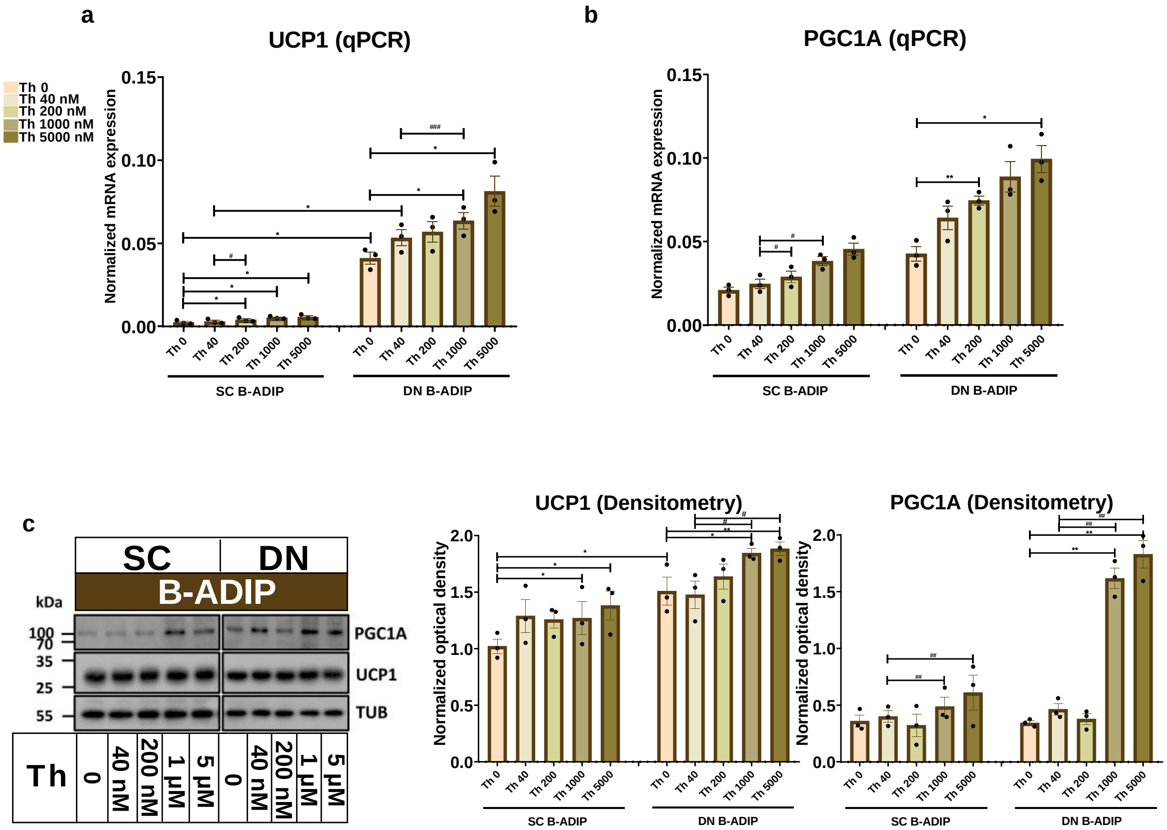


**Supplementary Figure 2.** Effect of gradually increasing concentrations of thiamine (Th) on thermogenic gene and protein expression in human subcutaneous (SC) and deep neck (DN)-derived brown differentiated adipocytes (B-ADIPs). (a-b) mRNA expression of *UCP1* and *PGC1a* assessed by RT-qPCR, n=3. (c) UCP1 and PGC1a protein expression detected by immunoblotting, n=3. Statistical analysis was performed by one-way ANOVA, *#p<0.05, **##p<0.01, ***###p<0.001, *comparing data at each concentration of Th to the lack of Th or # comparing the indicated groups.

a

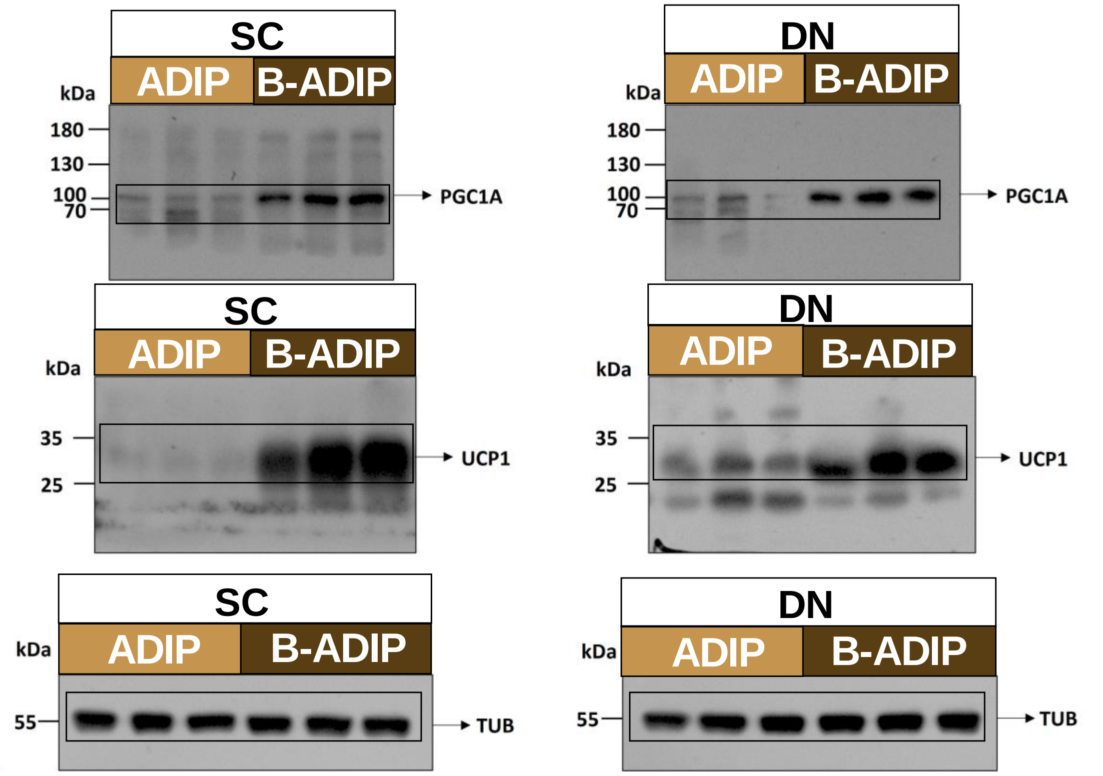
SC
ADIP
B-ADIP
kDa
180
130
100
70
PGC1A
DN
ADIP
B-ADIP
kDa
180
130
100
70
PGC1A
SC
ADIP
B-ADIP
kDa
35
25
UCP1
DN
ADIP
B-ADIP
kDa
35
25
UCP1
SC
ADIP
B-ADIP
kDa
55
TUB
DN
ADIP
B-ADIP
kDa
55
TUB

b

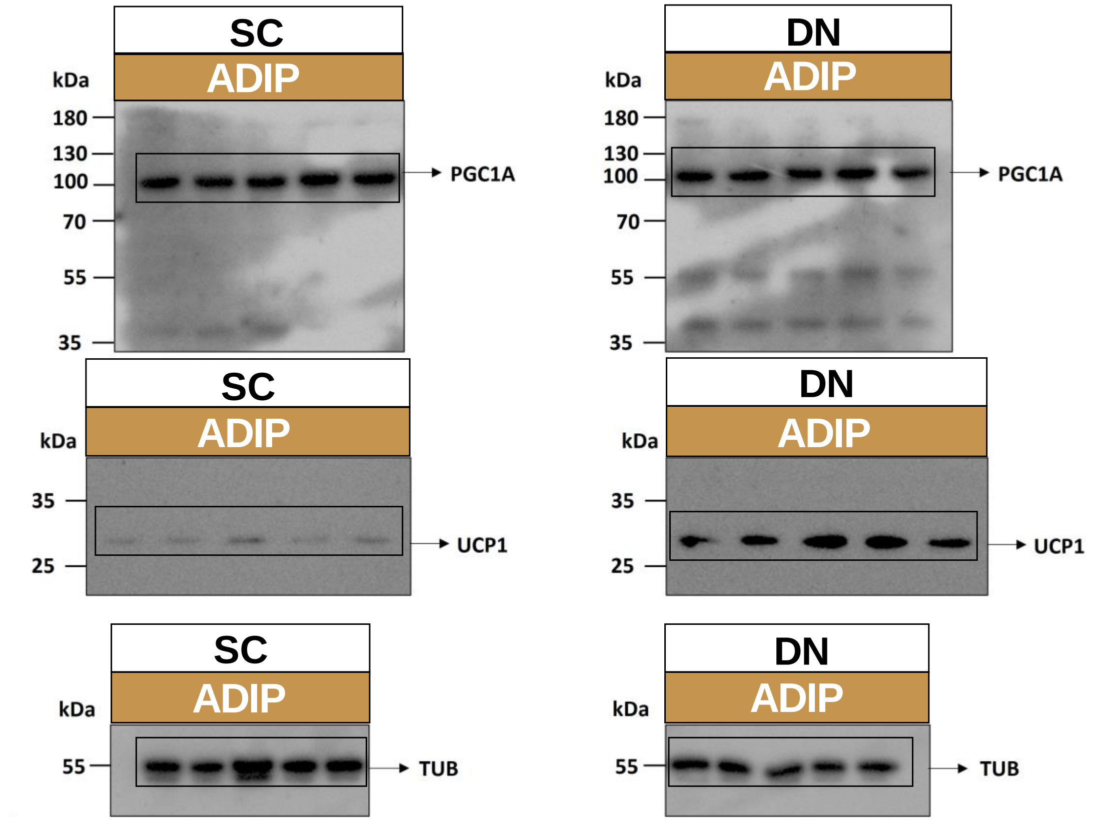
SC
ADIP
kDa
180
130
100
70
55
35
PGC1A
DN
ADIP
kDa
180
130
100
70
55
35
PGC1A
SC
ADIP
kDa
35
25
UCP1
DN
ADIP
kDa
35
25
UCP1
SC
ADIP
kDa
55
TUB
DN
ADIP
kDa
55
TUB

**Supplementary Figure 3.** Uncropped western blot images presented with molecular weight ladders, using MAB6158 monoclonal anti-UCP1 antibody or G0522 monoclonal anti-PGC1A antibody as shown in Figure 3c (a) and Figure 6c (b). Tubulin was used as endogenous control. Cropped areas are shown in black box regions.

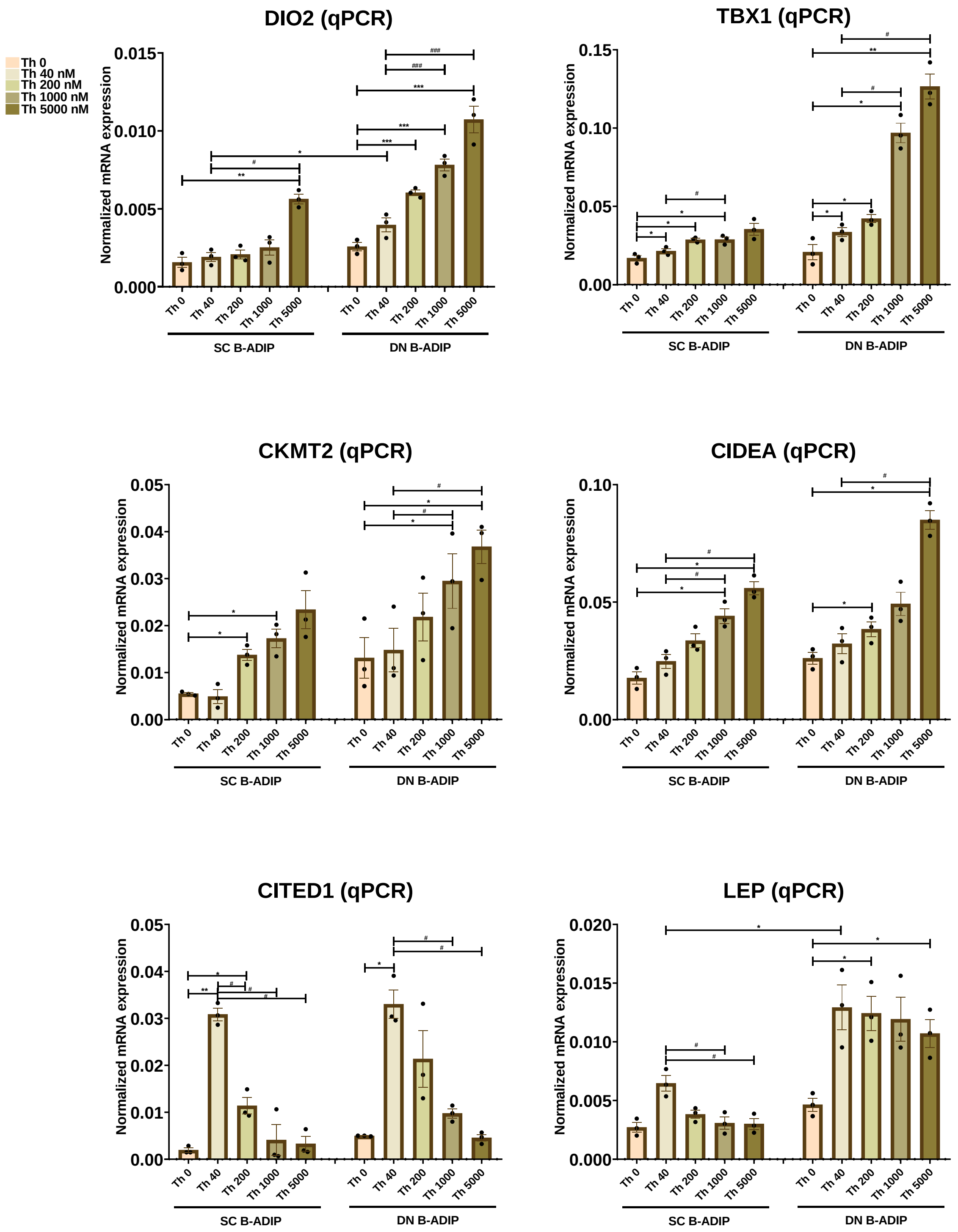

**Supplementary Figure 4.** Effect of gradually increasing concentrations of thiamine (Th) on thermogenic gene induction in human subcutaneous (SC) and deep neck (DN)-derived brown differentiated adipocytes (B-ADIPs). mRNA expression of *DIO2*, *TBX1*, *CKMT2*, *CIDEA*, *CITED1*, and *LEP* assessed by RT-qPCR, n=3. Statistical analysis was performed by one-way ANOVA, *#p<0.05, **##p<0.01, ***###p<0.001, *comparing data at each concentration of Th to the lack of Th (Th 0) or # comparing the indicated groups.

### 1.2 Supplementary Tables

**Supplementary Table 1.** Gene primers and probes

| GENES | ASSAY ID |
|---|---|
| *CIDEA* | Hs00154455_m1 |
| *CITED1* | Hs00918445_g1 |
| *CKMT2* | Hs00176502_m1 |
| *DIO2* | Hs00255341_m1 |
| *GAPDH* | Hs99999905_m1 |
| *LEP* | Hs00174877_m1 |
| *PPARGC1A* | Hs01016719_m1 |
| *SLC19A2* | Hs00949693_m1 |
| *SLC19A3* | Hs00228858_m1 |
| *SLC25A19* | Hs01001439_m1 |
| *TBX1* | Hs00271949_m1 |
| *UCP1* | Hs00222453_m1 |

**Supplementary Table 2.** Antibodies used in immunoblotting

| ANTIBODY | COMPANY | CATALOG NUMBER | DILUTION |
|---|---|---|---|
| UCP1 | R&D Systems, Minneapolis, MN, USA | MAB6158 | 1:750 |
| SLC19A3 | Novus Biologicals, Centennial, CO, USA | NBP1-69703 | 1:500 |
| SLC19A2 | Abcam, Cambridge, MA, USA | Ab229680 | 1:500 |
| PGC1α | Novus Biologicals, Centennial, CO, USA | NBP1-04676 | 1:1000 |
| Total OXPHOS | Abcam, Cambridge, MA, USA | ab110411 | 1:1000 |
| TUBULIN | Santa Cruz, USA | sc-5274 | 1:10000 |
| HRP-conjugated goat anti-rabbit IgG | Advansta, San Jose, CA, USA | R-05072-500 | 1:5000 |
| HRP-conjugated goat anti-mouse IgG | Advansta, San Jose, CA, USA | R-05071-500 | 1:5000 |
